# Association of the malate dehydrogenase-citrate synthase metabolon is modulated by intermediates of the Krebs tricarboxylic acid cycle

**DOI:** 10.1101/2021.08.06.455447

**Authors:** Joy Omini, Izabela Wojciechowska, Aleksandra Skirycz, Hideaki Moriyama, Toshihiro Obata

## Abstract

Mitochondrial malate dehydrogenase (MDH)-citrate synthase (CS) multi-enzyme complex is a part of the Krebs tricarboxylic acid (TCA) cycle ‘metabolon’ which is enzyme machinery catalyzing sequential reactions without diffusion of reaction intermediates into a bulk matrix. This complex is assumed to be a dynamic structure involved in the regulation of the cycle by enhancing metabolic flux. Microscale Thermophoresis analysis of the porcine heart MDH-CS complex revealed that substrates of the MDH and CS reactions, NAD^+^ and acetyl-CoA, enhance complex association while products of the reactions, NADH and citrate, weaken the affinity of the complex. Oxaloacetate enhanced the interaction only when it was presented together with acetyl-CoA. Structural modeling using published CS structures suggested that the binding of these substrates can stabilize the closed format of CS which favors the MDH-CS association. Two other TCA cycle intermediates, ATP, and low pH also enhanced the association of the complex. These results suggest that dynamic formation of the MDH-CS multi-enzyme complex is modulated by metabolic factors responding to respiratory metabolism, and it may function in the feedback regulation of the cycle and adjacent metabolic pathways.

## Introduction

Multi-enzyme complexes are involved in various metabolic pathways in the cell^1^, including the Krebs-tricarboxylic acid (TCA) cycle where citrate synthase (CS) and malate dehydrogenase (MDH) form a complex by an electrostatic interaction involving arginine residues of the CS (65Arg and 67Arg)^2^. This multi-enzyme complex channels the reaction intermediate, oxalacetate (OAA), within the complex to prevent its diffusion to the cellular matrix^2,3^. The formation of such a multi-enzyme complex mediating substrate channeling, so-called ‘metabolon’^4^, has many potential advantages including enhancement of pathway flux due to increased local substrate concentration near the enzyme reaction center, protection of intermediates and catalytic centers from inhibitors and competing pathways, and increased stability of intermediates^3,5,6^. The MDH-CS multi-enzyme complex is particularly interesting because concentrations of OAA in the mitochondrial matrix are believed to be insufficient to sustain the determined rate of the TCA cycle^7^. MDH forward reaction producing OAA is thermodynamically unfavorable at physiological concentrations of OAA^8^, hence the MDH-CS complex and metabolite channeling has been considered essential to achieve high OAA concentration at the CS reaction center to overcome the thermodynamic limitations^5,8,9^.

The TCA cycle feeds anabolic processes and participates in energy metabolism, being responsible for the oxidation of respiratory substrates to drive ATP synthesis. Oxidative phosphorylation involves the pumping of protons by the respiratory chain and back-flux of protons across ATP synthase leading to changes in matrix pH^10^. The TCA cycle is very tightly regulated by various factors in the mitochondrial matrix^5^. The cycle is regulated primarily by the mitochondrial NAD^+^/NADH ratio which is directly related to the mitochondrial ATP/ADP ratio that signals energy level in the cell^11^. NADH inhibits all the regulatory enzymes of the cycle, it functions as a feedback inhibitor to downregulate the TCA cycle and further the ATP production^5,11^. Additionally, ample acetyl-CoA (which is a product of oxidative pathways including glycolysis and lipid β-oxidation) upregulates flux through the cycle as an allosteric regulator causing a conformational change to the CS structure to enhance catalytic activity^12^. Thus, activities of the TCA cycle enzymes are regulated by various products and substrates of oxidative pathways and anabolic processes. Therefore, it is reasonable to assume that these allosteric regulators control not only the enzyme activities but also transient multi-enzyme complex formation through conformational changes of individual enzymes. We hypothesize that metabolites whose levels fluctuate in the mitochondrial matrix depending on the metabolic flux through the TCA cycle and adjacent metabolic pathways regulate multi-enzyme complex formation as a means of feedback regulation of these pathways. In this work, we explore the effects of metabolites and pH on the affinity of the MDH-CS multi-enzyme complex *in vitro*. We also computationally evaluate the effect of ligand binding on already established CS structures and accessibility of the 65Arg and 67Arg residues for multi-enzyme complex formation. The results showed the effects of the metabolic intermediates on the affinity of the MDH-CS complex, suggestive of the dynamics and function of the multi-enzyme complex in the regulation of the TCA cycle and adjacent pathways.

## Results

To determine the effect on the binding affinity of the MDH-CS complex, we assessed the affinity of the complex in the presence of various metabolites in the mitochondrial matrix whose concentrations are expected to alter depending on the respiratory activity. The *K*_d_ value of MDH-CS interaction was determined by MST as 2.29 ± 0.46 μM in the control condition when CS was used as the ligand (Fig. 1A, green curve). No binding was observed when MDH was incubated with bovine serum albumin (Fig. S1 in Supplementary Information, orange curve).

**Figure 1.**
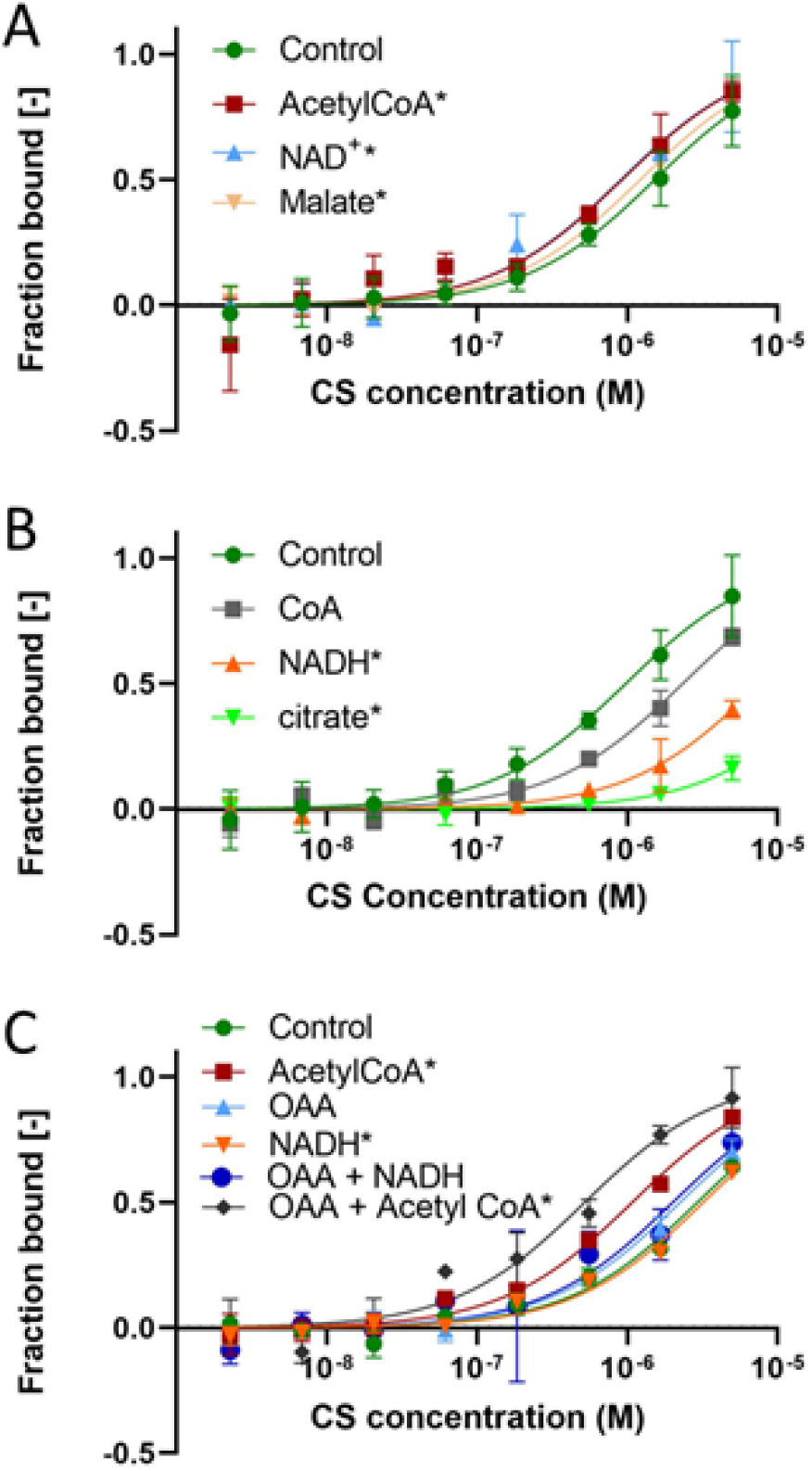
Effects of metabolites involved in the MDH and CS reactions on the affinity of the MDH-CS multi-enzyme complex. The affinity of the MDH-CS multi-enzyme complex was analyzed by microscale thermophoresis (MST) using fluorescently labeled MDH and CS as the ligand. Curves represent the response (fraction bound) against CS concentration (M). The interaction was assessed in the MST buffer (control, green) or those with 10 mM of metabolites. Error bars represent the standard deviations of three measurements. Asterisks indicate the conditions that showed significant *K*_d_ differences with no *K*_d_ confidence overlap with the control. **(A)** Effects of reaction substrates of MDH and CS. The MDH-CS interaction was assessed in the presence of acetyl-CoA (red), NAD^+^ (blue), or malate (brown). **(B)** Effects of reaction products of CS and MDH. The MDH-CS interaction was assessed in the presence of CoA (grey), NADH (orange), or citrate (green). **(C)** Effects of the oxalacetate (OAA) in combination with other CS substrates. The effects of sole substrates, acetyl-CoA (red square), OAA (blue), and NADH (orange), as well as their combinations, OAA/acetyl-CoA (purple) and OAA/NADH (gray), were analyzed.

### Effects of the metabolites involved in the reactions catalyzed by MDH and CS

The presence of 10 mM of the substrates of MDH and CS reactions led to significant changes in the *K*_d_ of the complex. The addition of acetyl-CoA and NAD^+^ significantly reduced the *K*_d_ to 1.10 ± 0.23 and 0.89 ± 0.26, while malate slightly but not significantly reduced *K_d_* to 1.80 ± 0.17 μM (Fig. 1A). On the other hand, products of these reactions, NADH and citrate, significantly increased the *K*_d_ values to 7.50 ± 0.60 μM and 25.1 ± 6.3 μM, respectively (Fig. 1B). The effect of CoA on the affinity of this complex was not significant (*K*_d_ 2.32 ± 1.15 μM; Fig. 1B). Thus, substrates of the MDH and CS reactions increased complex association while products of the reaction decreased it.

MDH-CS multienzyme complex channels oxaloacetate by electropositive interaction^2,13^. Oxaloacetate did not alter the dissociation constant of this complex (*K*_d_ = 2.23 ± 0.60 μM; Fig. 1C). We evaluated the effect of oxaloacetate, in the presence of NADH or acetyl-CoA which are other substrates involved in the enzyme reactions. The addition of NADH together with oxaloacetate (Fig. 1C) did not significantly alter the binding affinity of MDH and CS (*K*_d_ = 2.01 ± 0.79 μM). However, oxaloacetate together with acetyl-CoA significantly reduced the *K*_d_ to 0.52 ± 0.11 μM (Fig. 1C) in comparison to those at both the control condition and the presence of acetyl-CoA. Hence, OAA altered the affinity of the multienzyme complex association only in the presence of acetyl-CoA.

### Alteration of CS structure by the interaction with acetyl-CoA and OAA

To gain insight into how two substrates of the CS reaction synergistically increased the MDH-CS binding affinity, we evaluated the effect of these intermediates on the structure and conformation of CS. Two previously established porcine CS structures, which were solved in the presence of citrate and citrate with CoA referred to as the open (PDB 1cts; Fig. 2A) and closed formats (PDB 2cts; Fig. 2C), respectively^14^, were used for computational analyses. These formats are believed to be the form to accept the substrates and release the products and that to carry out catalytic steps with its active site buried deep within the protein, respectively^15^. The bottom half domain was well conserved, and the top half was inconsistent between these two conformations (Fig. 2B). The distribution of the surface charges was reallocated between the open and closed formats (Fig. 2D and E). While the patches are negatively charged and hydrophobic on the surface near the active site in the open format (Fig. 2D), positively charged patches become dominant in the closed format (Fig. 2E). Acetyl-CoA forms hydrogen bonds with the 274His and 375Asp at the CS active site as the first step in the reaction^16^. The structural modeling of the acetyl-CoA localization in the closed format of CS suggested that acetyl-CoA also forms hydrogen bonds with A46Arg and B164Arg (A and B represent the subunits of the CS homodimer) in the bottom conserved domain, and B164Arg is connected to A366Lys in the top mobile domain through acetyl-CoA (Fig. 2B). Thus, the presence of acetyl-CoA in the catalytic site of CS likely favors the closed format of CS protein by forming a hydrogen bond network.

**Figure 2.**
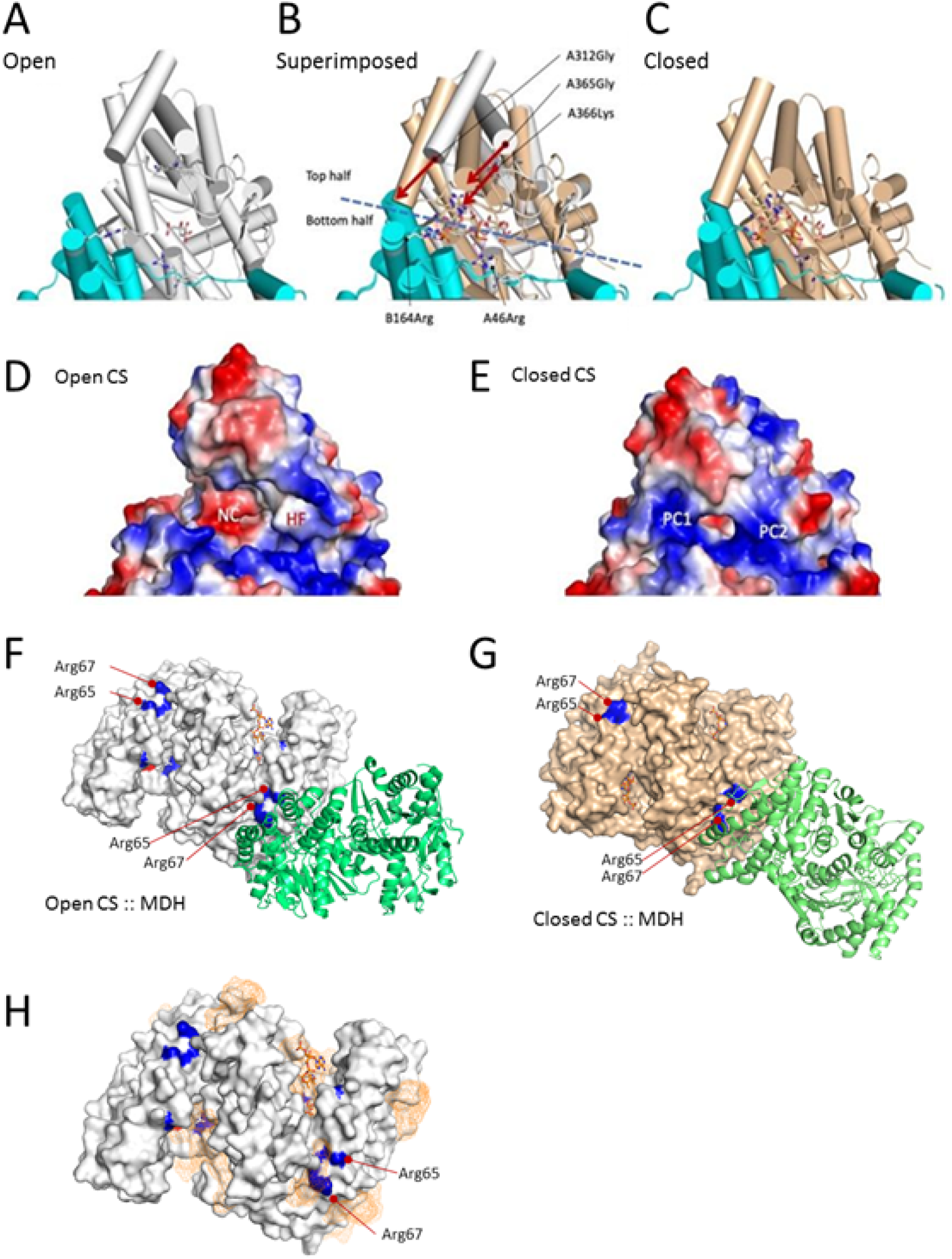
Effect of acetyl-CoA binding on CS structure. **(A)** The open format of CS (PDB ID, 1cts) in a cartoon model, with cylindrical α-helices containing a citrate molecule in a stick model. Subunits A and B are colored white and cyan, respectively. Key residues, A266Lys, A46Arg, and B164Arg, are shown in the sticks’ side chains. The residues are shown in the order of the chain name, residue number, and amino acid. the dimer structure was generated using crystallographic symmetry. One active site domain composed mainly by the A-chain is shown. **(B)** The superimposed model between the open (1cts; white) and close (2cts; wheat) formats. The citrate molecule is at the same location with a slight rotation. The CoA molecule is present only in the closed format. The molecular domain shown can be divided into the movable upper half and the rigid bottom half. In the bottom domain, A45Arg and B146Arg are shown in the stick model. The locations of those two Cα in Arg are almost consistent between the open and close formats. The top half domain is movable. The motion is visible as the rotation of α- helices represented by A312Gly and A365Gly, indicated by arrows. The A366Lys moves inward and forms a hydrogen bond network A366Lys (NZ):: A438COA(O8A):: B164Arg(NH1). **(C)** The closed format of CS (PDB ID, 2cts) with citrate and CoA molecules in stick models. **(D)** The surface electrostatic potential of the open format CS excluding ligands. Calculations were performed in the vacuum environment and ranged between −71 and +71. Red and blue represent the negative and positive potentials. The domain shown corresponds to panel A. Patches of negative charge (NC) and hydrophobic area (HF) are observed. **(E)** The surface electrostatic potential of the closed format CS excluding ligands. The domain shown corresponds to panel D. Patches of positive charge (PC1, PC2) are observed. **(F and G)** Simulated interactions between MDH (green) and CS in open (F; white) and close (G; wheat) formats. The 65Arg and 67Arg residues that are involved in the MDH-CS interaction are highlighted in blue. Active site residues, His 274, His320 (blue), and Asp375 (red) were shown. Residues involved in the binding are listed in Table S2. **(H)** Predicted acetyl-CoA binding sites in CS apoenzyme. The white surface model of the CS apoenzyme in the open format is shown. The 274His, 320His, 65Arg, and 67Arg residues are highlighted in blue. The orange mesh indicates the positions of acetyl-CoA at the predicted binding sites. White and orange stick models indicate the citrate and CoA in the reported crystal structure, respectively.

MDH protein (PDB 1mld) is predicted to interact with open and closed CS differently (Fig. 2F&G). ZDOC score, a relative binding energy function calculated by ZDOC, of the 65Arg residue was larger in closed format than open format, indicating that CS in closed format binds better with MDH (Table S2 in Supplementary Information). The closed CS also has a larger accessible surface area, buried surface area after binding, and more hydrogen bonds with MDH than the open format (Table S2 in Supplementary Information). These results indicate that the CS in the closed format favors MDH binding.

Acetyl-CoA also allosterically regulates CS activity^17^. Our structural modeling predicted one potential acetyl-CoA binding site in the CS apoenzyme in the open format that locates very close to the 65Arg and 67Arg involved in the interaction between MDH and CS (Fig. 2F).

### Effects of other TCA cycle intermediates

Since the TCA cycle intermediates are known to allosterically regulate the activities of the enzymes of the cycle, we tested the effect of these intermediates on the binding affinity of the MDH-CS multi-enzyme complex. Succinate and α-ketoglutarate significantly reduced the *K*_d_ of the interaction to 0.18 ± 0.41 μM and 1.14 ± 0.21 μM, respectively (Table 1). No significant effect on the MDH-CS complex affinity was observed by the addition of fumarate and succinyl-CoA into the test solution (Table 1). Thus succinate and α-ketoglutarate increased the binding affinity of this complex, while citrate decreased binding affinity (Table 1, Fig. 1B).

**Table 1.** Effects of the TCA cycle intermediates on the affinity of the MDH-CS complex

| Intermediate (10mM) | $K_d \pm K_d$ confidence ($\mu\text{M}$) |
| --- | --- |
| control | $2.29 \pm 0.46$ |
| citrate | $25.1 \pm 6.3^*$ |
| oxaloacetate | $2.23 \pm 0.60$ |
| malate | $1.85 \pm 0.17$ |
| succinyl CoA | $1.97 \pm 0.48$ |
| α-ketoglutarate | $1.14 \pm 0.21^*$ |
| fumarate | $1.58 \pm 0.37$ |
| succinate | $0.18 \pm 0.41^*$ |

Data are presented as $K_d \pm K_d$ confidence. Asterisks represent metabolites that showed significant differences with no $K_d$ confidence overlap with the control. The data for citrate, oxaloacetate, and malate are the same as Fig. 1.

### Effects of the indicators of mitochondrial energy status

The TCA cycle is an amphibolic pathway. The catabolic role of this cycle is to generate reducing equivalents that are used in the electron transport chain for oxidative phosphorylation. The energy level in the cell is closely related to respiratory activities which must be constantly regulated in response to metabolic conditions. To determine whether energy molecules in the mitochondria have any effect on the binding affinity of the MDH-CS complex, we evaluated the binding affinity of the complex in the presence of 10 mM AMP, ADP, and ATP. AMP and ADP did not significantly affect the *K*_d_ values (3.10 ± 0.62 μM and 2.24 ± 0.56 μM for AMP and ADP, respectively; Fig. 3A). There was a significant increase in the binding affinity of the MDH-CS complex in the presence of ATP with the *K*_d_ reduced to 0.46 ± 0.03 μM (Fig. 3A). The reduction of *K*_d_ was also observed at lower concentrations of ATP with a dose-dependent trend; the *K*_d_ value reduced with increasing ATP concentration (Fig. 3B). The presence of 1.25, 5, 10, and 20 mM of ATP significantly reduced the *K*_d_ of MDH-CS interaction to 1.06 ± 0.26, 0.77 ± 0.30, 0.76 ± 0.11, and 0.72 ± 0.15 μM, respectively, in comparison to the control condition. The pH in the mitochondrial matrix changes in response to the electron transport chain activities which are closely related to the energy status^10^. Resting matrix pH ranges between pH 7 and 8, and when protons diffuse back into the matrix to drive ATP synthesis, the matrix pH becomes more acidic down to pH 6^18^. To assess the effect of pH in the mitochondrial matrix on the MDH-CS complex affinity, we monitored binding affinity in the pH range between 6.0 and 8.0. MDH-CS interaction in pH 8 MST buffer (control) yielded a *K*_d_ of 2.29 ± 0.46 μM (Fig. 3C). At pH 6 and 7, the *K*_d_ values were 0.68 ± 0.05 μM and 1.60 ± 0.45 μM, respectively, and significant differences were observed when the control group (pH 8) was compared with pH 6.

**Fig. 3.**
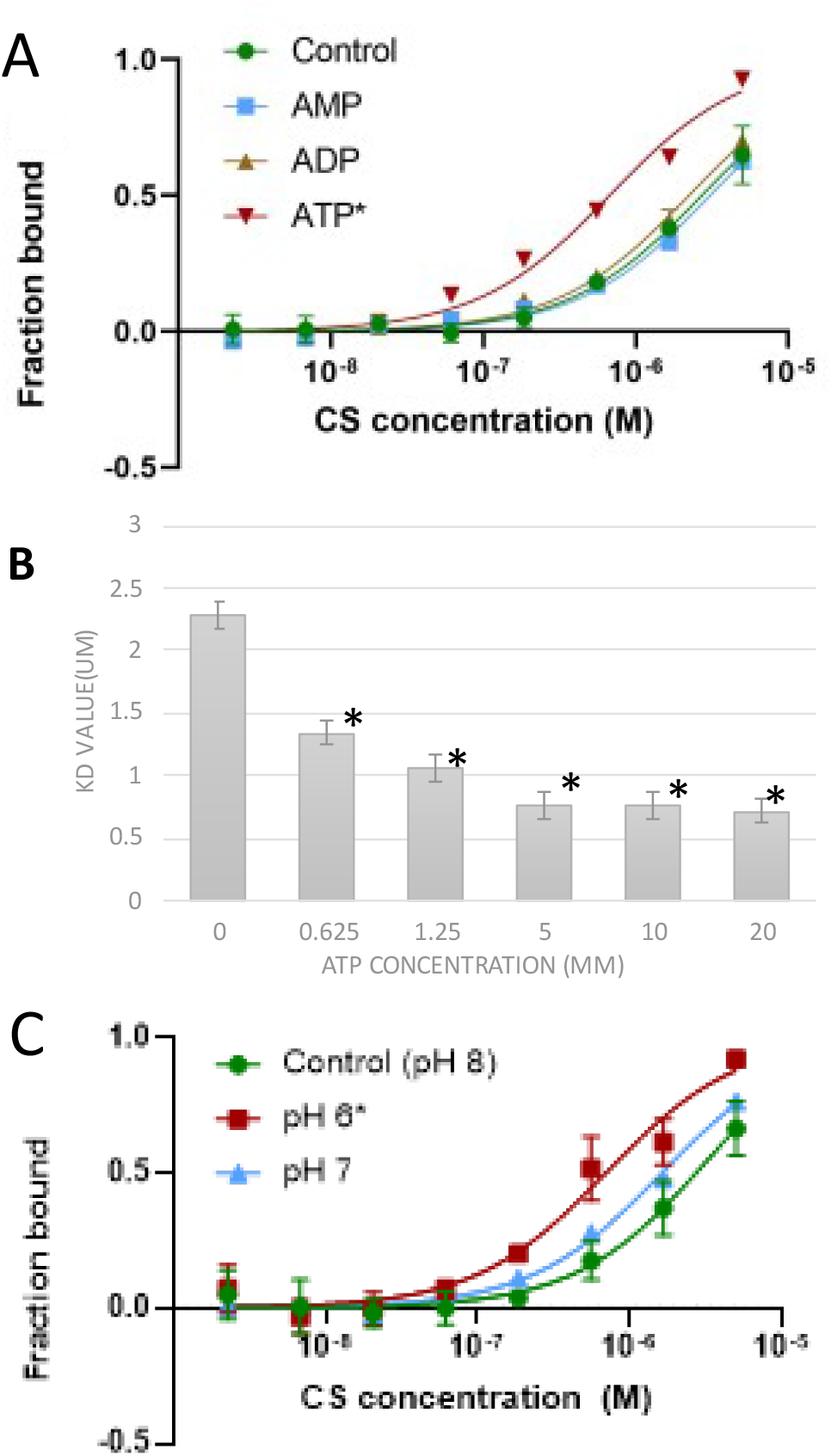
Effects of mitochondrial energy status on the affinity of the MDH-CS multi-enzyme complex. The affinity of the MDH-CS multi-enzyme complex was analyzed by microscale thermophoresis (MST) using fluorescently labeled MDH and CS as the ligand. Points and bars represent the means, and the error bars represent the standard deviations of three measurements. Asterisks indicate the conditions that showed significant *K*_d_ differences with no *K*_d_ confidence overlap with the control. **(A)** Effects of 10 mM ATP (red), ADP (brown), and AMP (blue) on the interaction between CS and MDH. Curves represent the response (fraction bound) against CS concentration (M). The interaction was assessed in the MST buffer (control, green circle) or those with 10 mM of metabolites. **(B)** Dose-dependent effect of ATP on the dissociation constant (*K*_d_) of the MDH-CS interaction. *K*_d_ was calculated from the response curve against CS concentration. **(C)** Effects of pH. Curves represent the response (fraction bound) against CS concentration. The interaction was assessed in the MST buffer with pH 8.0 (control, green) or that with pH 6.0 (red) and 7.0 (blue).

## Discussion

Here, we report the effects of metabolic intermediates as regulators of MDH-CS multi-enzyme complex formation *in vitro*. Many of the tested intermediates related to the TCA cycle either enhanced or reduced the affinity of the enzyme complex, suggesting their functions in the dynamic regulation of the TCA cycle metabolon in response to the metabolic status in the mitochondrial matrix. We tested the effects of metabolites at a high concentration of 10 mM to capture any possible effects of compounds. This is a much higher concentration than the reported concentrations of the metabolites in the mitochondria^19^, which is calculated with an assumption of equal metabolite distribution in the mitochondria. However, a higher concentration is achievable by local metabolite accumulation in microcompartments such as a metabolon^20–22^. It should be noted that the effects of these metabolites on the MDH-CS complex are compound-specific as some metabolites showed no effect even at this high concentration. Additionally, ATP had significant effects on MDH-CS binding even at the reported concentration range in mammalian mitochondria^19^. Thus, the observed effects of metabolites on MDH-CS interaction are likely reproducible in living cells.

Interestingly, the compounds produced by the MDH and CS reactions (citrate and NADH) reduced the affinity of the complex, whereas the substrates of the reactions (acetyl-CoA and NAD^+^) and the TCA cycle intermediates (α- ketoglutarate and succinate) enhanced it. These results suggest the possible function of the MDH-CS multi-enzyme complex in feedback regulation of the TCA cycle flux. Assuming that the metabolon enhances the MDH-CS forward reactions, the responses of the MDH-CS complex to the metabolite availabilities would facilitate the forward flux of the cycle when the substrates of the reactions are abundant and suppress it when the products accumulate. The majority of the TCA cycle enzymes are known to be allosterically regulated by various compounds related to the cycle^5^. Our results indicate that the allosteric regulation also affects the metabolon formation and further contributes to the fine-tuning of the TCA cycle flux.

It should be noted that the metabolites that affected the MDH-CS complex affinity in this study are also involved in other metabolic pathways. Therefore, the observed effects of the intermediates may also function in balancing metabolic fluxes through the TCA cycle and the related pathways. For instance, citrate can function to balance the metabolic flux through the TCA cycle and fatty acid biosynthesis. The results in this study indicate that citrate may downregulate the CS reaction by the inhibition of MDH-CS interaction. Accumulation of citrate also likely stimulates mitochondrial fatty acid synthesis by promoting the polymerization of acetyl-CoA carboxylase to activate it^23^. α-ketoglutarate and succinate enhanced the interaction of the MDH-CS multi-enzyme complex in this study. While these metabolites are TCA cycle intermediates, α-ketoglutarate is also derived from the transamination of amino acids^24^. Succinate is a substrate for succinate dehydrogenase of the electron transport chain (complex II), which couples the TCA cycle and the respiratory chain^25^. These metabolites both enter the TCA cycle as anaplerotic intermediates. These oxidative pathways are activated to produce ATP when the energy status is low^26^. Therefore, it is reasonable that these metabolites facilitate the MDH-CS interaction to enhance the TCA cycle flux to further enhance the production of reducing equivalent and ATP. OAA is another metabolite at a branching point of metabolic pathways. OAA is a substrate for aspartate aminotransferase (AAT) which converts OAA to aspartate using glutamate as the amino donor. As the AAT reaction mediates significantly higher flux than the TCA cycle at least in some mammalian tissues^27^, the TCA cycle competes for OAA with the AAT reaction in vivo. Accumulation of acetyl-CoA likely facilitates metabolic flux through the TCA cycle over the AAT pathway at this branch point by promoting MDH-CS complex formation and metabolite channeling of OAA to limit its accessibility to AAT^9^. Interestingly, the co-presence of both acetyl-CoA and OAA further enhances the interaction in this study, which most likely strengthens the re-direction of metabolic flux to the TCA cycle. Such a multi-step mechanism may enable the fine-tuning of the metabolic flux through the pathways.

The effects of CS substrates on the MDH-CS complex affinity observed in this study are partially explained by the conformational changes taking place when compounds are bound and released from the enzymes. The binding of acetyl-CoA to the substrate-binding site puts CS in the closed format which favors MDH binding. Additionally, dominant positive charges in the closed format likely enhance electrostatic interaction with MDH. This positive electrostatic potential extending over much of the surface of the protein probably also favors metabolite channeling by the electrostatic interactions with the negatively charged substrate^28,29^. These suggest that CS is in the closed format in the MDH-CS metabolon, while direct experimental evidence is missing.

OAA alone did not affect the binding affinity of the two enzymes. This agrees with the report of Tompa et al.^30^ in which 10^−4^ M OAA did not affect the enzyme interaction. The insignificant effect of OAA might be due to the location of the OAA binding site deep in the cleft of the catalytic active site (Fig. 2) which would have a limited effect on the transition to the closed format. On the other hand, acetyl-CoA makes hydrogen bonds with residues in the mobile domain and can contribute to stabilizing the closed format. In the presence of acetyl-CoA, OAA significantly increased the affinity of the MDH-CS complex likely by further stabilizing the close format in which the CS reaction takes place. Alternatively, this effect could be a result of a conformational change that occurs when acetyl-CoA binds to the CS active site. Acetyl-CoA is considered important for the proper positioning of OAA in the active site of CS. When acetyl-CoA is absent, CS is highly flexible and unstructured, which prevents OAA from proper positioning in the active site^31^. These can explain the need for acetyl-CoA in the function of OAA to modulate the interaction of the MDH-CS complex.

The structural modeling in this study also indicates that the binding of acetyl-CoA to potential allosteric sites can facilitate MDH-CS complex formation by bringing MDH residues closer to the CS arginine residues which contribute to the electrostatic forces responsible for the protein-protein interaction^2^. We observed that α-ketoglutarate, which has been reported as allosteric effectors of the CS activity^32–34^ also significantly increased the binding affinity of the MDH-CS complex. Our simulation indicated that this compound did not share the same binding site as acetyl-CoA (Fig. S2 in Supplementary Information), indicating that the enhancing effect of ATP and α-ketoglutarate on the multi-enzyme complex formation is likely not based on the same mechanism as that of acetyl-CoA.

Mitochondrial energy status can also affect the formation of the MDH-CS complex. It is reasonable since energy production is one of the significant outcomes of respiratory metabolism. ATP plays an important role in the regulation of metabolism and it fluctuates with changes in metabolic conditions. It has been reported as an allosteric^33^ and competitive inhibitor of CS for acetyl-CoA^32^. In this study, we observed a concentration-dependent positive effect of ATP on the binding affinity of the MDH-CS complex. Variations in the pH of the mitochondrial matrix reflect the pumping of protons by the respiratory chain and back-flux of protons across the ATP synthase^10^ and range between pH 6 to 8^35,36^. Protons return to the matrix ‘through’ ATP synthase driving the synthesis of ATP^37^, and an increase in H^+^ concentration will lead to a slight drop in pH. The significant difference in binding affinity of MDH and CS was observed when the pH 6 condition was compared to pH 8, suggesting that the MDH-CS complex formation is likely affected by the pH changes in the mitochondrial matrix that occur with ATP synthesis. Our results indicate that the energy-rich conditions can favor the interaction of MDH-CS, which is expected to further increase ATP production, although the metabolic and physiological relevance of this regulation is still unclear.

Our study showed that substrates of individual MDH and CS enzymes, which have been reported to increase the reaction rates, favor MDH-CS multi-enzyme complex association. On the other hand, reaction products that inhibit the forward reactions favor dissociation of the complex. The results also suggest that metabolites used for anabolic processes dissociate the complex while products of catabolic processes enhance it. These results indicate that the TCA cycle metabolon would dynamically interact depending on the accumulation of metabolic intermediates and is probably involved in a switch between catabolic and anabolic processes as a mechanism underlying metabolic network regulation. The dynamics of the MDH-CS complex association and its relationship with metabolic flux distributions in central carbon metabolism and local metabolite concentrations in the mitochondrial matrix should be analyzed in a living system to evaluate the metabolic functions of the dynamic metabolon.

## Methods

### Materials

Mitochondrial CS (#SLCC7842) and MDH (#SLCD5169) isolated from porcine hearts were purchased from Millipore Sigma (München, Germany). MST dilution Buffer was composed of 0.1 M potassium phosphate buffer containing 0.05% Tween 20 with pH 8 unless otherwise stated.

### Sample Preparation

Fluorescence labeling of the target protein (MDH) was performed by Red N-hydroxy succinimide (NHS) 2nd generation (NanoTemper Technologies GmbH, München, Germany) following the manufacturer’s protocol. Ammonium sulfate suspension of the MDH was centrifuged at 23,000 × *g* for 1 min at 4 °C and the precipitate was resuspended in the labeling buffer to gain 90 μL of 10 μM MDH and mixed with 10 μL of 300 μM of dye (NanoTemper Technologies) prepared in dimethyl sulfoxide to create 100 μl of dye-protein solution. This was incubated in the dark for 30 min at room temperature. Unbound fluorophores were removed by a desalting column using MST buffer as the running buffer. Labeled protein was diluted with MST buffer to obtain final fluorescent counts between 500-1000 FU. Fifty microlitres of ammonium sulfate suspension of the ligand protein (CS) was centrifuged and resuspended in 25 μl of the MST buffer to gain 30 μM CS solution. A twofold CS dilution series was prepared with the MST buffer across eight tubes. A series of eight dilutions was used because there was no binding at lower concentrations, and higher concentrations of CS were difficult to be achieved with the commercially available enzyme products. No binding could be detected when CS was labeled and MDH was used as the ligand protein. Ten microliters of the diluted ligand protein solution were mixed with an equal volume of the target protein solution to be analyzed. The effects of the metabolites on the affinity of the MDH-CS complex were examined by adding the metabolites to the MST buffer to be 10 mM final concentrations. To test the effects of pH, the MST buffer was prepared with pH 6 and 7 potassium phosphate buffers.

### Microscale Thermophoresis (MST) Assay

MST experiments were performed by a Nano Temper Monolith NT.115 device (NanoTemper Technologies) using nano-red excitation. Samples were loaded into premium-coated capillaries (NanoTemper Technologies) and analyzed with MST power 40%, excitation power 80%, and the time windows of 5 sec before, 30 sec during, and 5 sec after the IR-laser. The temperature of the instrument was set to 23°C for all measurements.

### Data Analysis

All measurements were conducted with three replicates and the results were presented as mean values ± standard deviation. After recording the MST time traces, data were analyzed using the MO Affinity Analysis software (Nano Temper Technologies) by temperature jump (T-Jump) analysis, and the values obtained were normalized and plotted against the CS concentration. The dissociation constants (*K*_d_) with confidence intervals were determined using the *K*_d_ binding model. Graph Pad Prism 9.1.1 (GraphPad Software, San Diego, CA, USA) was used to create *K_d_*-binding curves. *K_d_* values were considered significantly different when there was no overlap of *K*_d_ intervals between the two groups.

### Structural Modeling

Structural mining and preparation of graphics were performed using PyMOL Molecular Graphics System, Version 2.4.1 (Schrödinger, LLC, New York, NY, USA). Both CS (PDB ID, 1cts and 2 cts), and MDH (1mld) structures were registered as asymmetric monomers. Dimer formats of enzymes were generated via the symmetry operations in PyMol. Apo format enzymes were subjected to the prediction of ligand binding on the Swiss Dock server^38^; http://www.swissdock.ch). Ligand structures were taken from the ZINC15 database ^39^ via the Swiss Dock server. Neighboring ligand sites were clustered by the Swiss Dock server (Table S1 in Supplementary Information). In the figures, only the first one structure in the first ten clusters are shown. Docking experiments were performed between MDH and either open or closed CS by the ZDOCK server^40^ (https://zdock.umassmed.edu). The binding was performed to fix the CS structures and rotate the MDH structure. Both CS and MDH were dimerized via symmetry operation using PyMOL. Complexes with the contact surface near 65Arg and 67Arg residues were selected from the top ten binding mates out of 2000 configurations. Surface areas and suffice interactions were evaluated using Proteins, Interfaces, Structures, and Assemblies (PDBePISA) server^41^ (https://www.ebi.ac.uk/pdbe/pisa/).

## Acknowledgment

This material is based upon work supported by the National Science Foundation Faculty Early Career Development Program (CAREER) to TO under Grant No. 1845451.

## Author contribution statement

T.O. conceived and designed the experiments; J.O., I.W., and A.S. established the MST procedure; J.O., I.W. conducted MST experiments; J.O. and T.O. analyzed the MST data; H.M. performed structural modeling; J.O. and T.O. wrote the manuscript; A.S. and H.M. edited the manuscript; all authors have approved the final version of the manuscript.

## Competing Interest Statement

The authors declare no competing interests.

## Supplementary Information

**Table S1.** Simulation results in ligand binding by Swiss Dock.

| Ligand | Total listed binding sites | Cluster | Energy in cluster 0 | Energy in cluster 9 |
| --- | --- | --- | --- | --- |
| Acetyl-CoA | 256 | 32 | −15.6943 | −29.8399 |
| α-ketoglutarate | 256 | 44 | −17.7629 | −4.75648 |
| succinyl-CoA | 256 | 32 | −35.1062 | −26.976 |
| ATP | 252 | 34 | −35.9538 | −30.2348 |

Energy (no dimension), relative energies provided by the Swiss Dock server. The greater the negative values, the stronger the binding.

**Table S2.** Properties of CS A subunit 65Arg-60Ser in complex with MDH A subunit.

| Parameters | Open (1cts) | Close (2cts) | Difference |
| --- | --- | --- | --- |
| ZDOCK Score (No. of selected structure) | 644.695 (4) | 742.992 (12) | +98.297 (+8) in 2cts |
| Accessible Surface Area, Å<sup>2</sup> | 284.54 | 325.35 | +40.81 in 2cts |
| Buried Surface Area before binding, Å<sup>2</sup> | 0.00 | 0.00 | 0.00 |
| Buried Surface Area after binding, Å<sup>2</sup> | 214.35 | 280.46 | +66.11 in 2cts |
| Hydrogen bonds | 4<br>## Structure 1 Dist. [Å] Structure 2<br>1 A:TYR 69[ N ] 3.51 A:ASP 88[ O ]<br>2 A:ARG 67[ NE ] 3.75 A:ASP 88[ OD1]<br>3 A:ARG 65[ NH2 ] 3.32 A:ASP 88[ OD2]<br>4 A:ASP 12[ OD1 ] 2.88 A:ASN 312[ ND2] | 6<br>## Structure 1 Dist. [Å] Structure 2<br>1 A:LYS 76[ NZ ] 2.60 A:VAL 249[ O ]<br>2 A:ARG 65[ NH1 ] 2.78 A:ASN 278[ O ]<br>3 A:ARG 20[ NH1 ] 2.82 A:GLU 289[ OE2]<br>4 A:GLU 104[ OE2 ] 3.55 A:ARG 167[ NH2]<br>5 A:TYR 69[ OH ] 3.53 A:CYS 188[ N ]<br>6 A:THR 103[ OG1 ] 3.71 A:VAL 192[ N ] | + 2 in 2cts |
| Salt bridges | 4<br>## Structure 1 Dist. [Å] Structure 2<br>1 A:ARG 67[ NE ] 3.75 A:ASP 88[ OD1]<br>2 A:ARG 67[ NH2 ] 3.73 A:ASP 88[ OD1]<br>3 A:ARG 65[ NH2 ] 3.32 A:ASP 88[ OD2]<br>4 A:LYS 16[ NZ ] 3.17 A:GLU 308[ OE1] | 4<br>## Structure 1 Dist. [Å] Structure 2<br>1 A:ARG 20[ NH1 ] 2.82 A:GLU 289[ OE2]<br>2 A:ARG 20[ NH2 ] 3.70 A:GLU 289[ OE2]<br>3 A:GLU 104[ OE2 ] 3.55 A:ARG 167[ NH2]<br>4 A:GLU 104[ OE2 ] 3.55 A:ARG 167[ NH2] | 0 |

**Figure S1.**
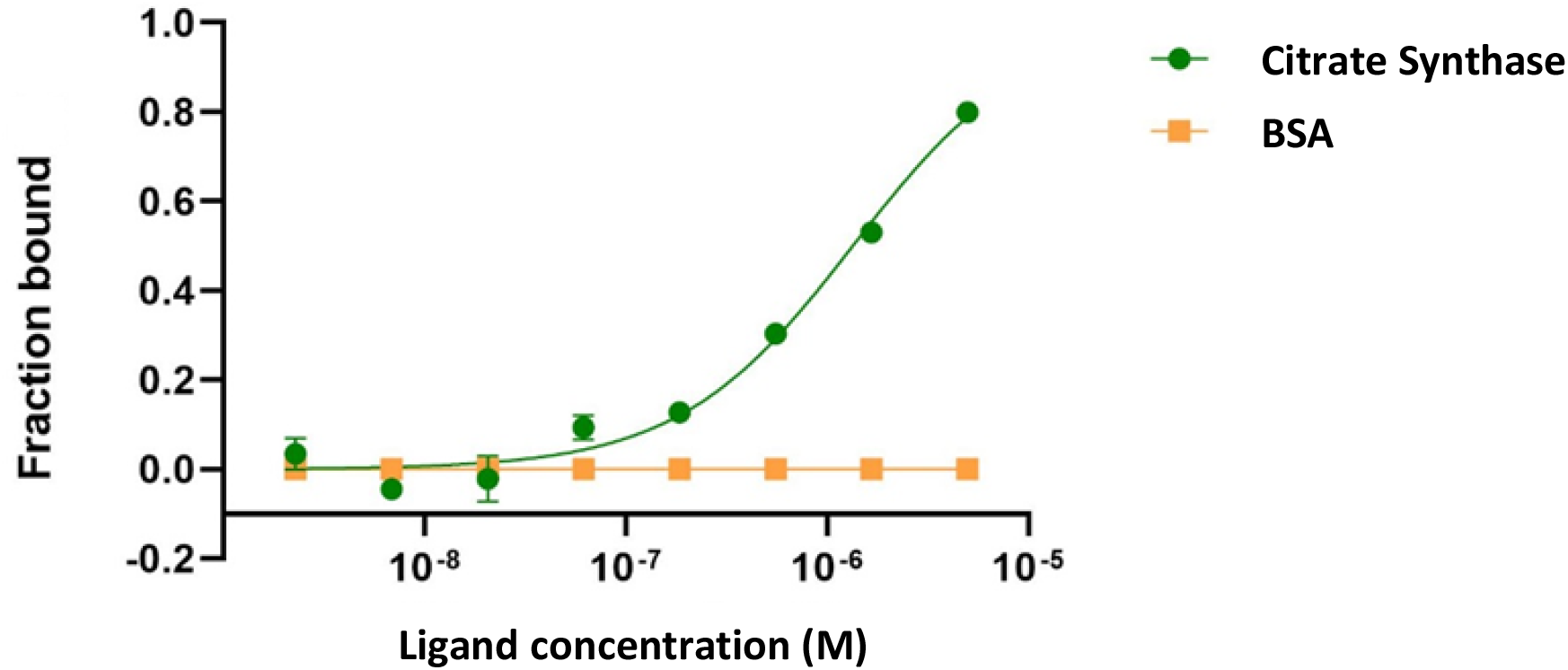
Microscale thermophoresis (MST) analysis of the MDH-CS multi-enzyme complex. The affinity of the MDH-CS multi-enzyme complex was analyzed by MST using fluorescently labeled MDH as the target. CS (green) and bovine serum albumin (orange) were used as the ligand proteins. Curves represent the response (fraction bound) against the concentrations (M) of ligand proteins. Error bars represent the standard deviations of three measurements.

**Figure S2.**
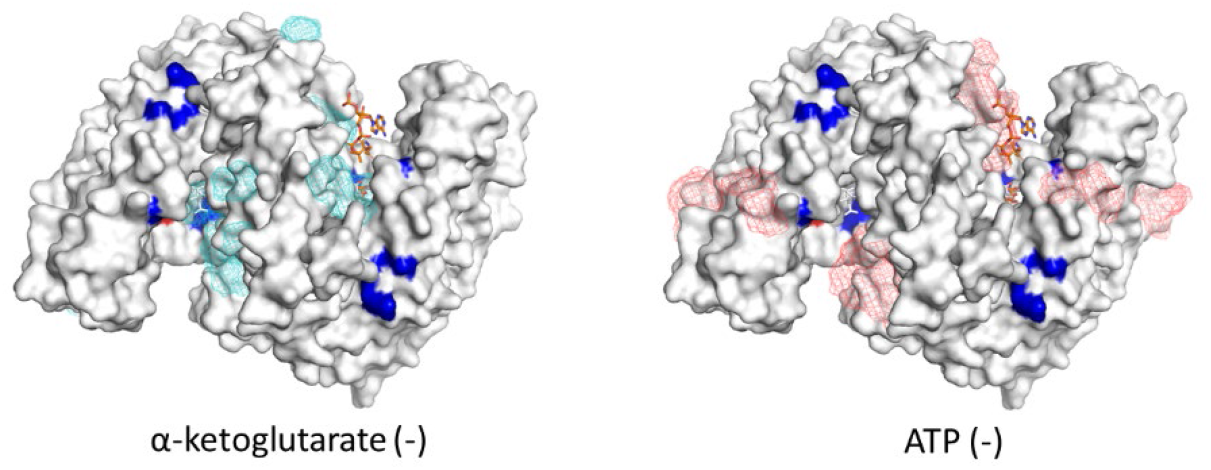
Predicted ligand binding sites in CS apoenzyme. The white surface models of the CS apoenzyme in the open format are shown. The 274His and 320 His residues at the reaction center and the 65Arg and 67Arg residues that are involved in the MDH-CS interaction are highlighted in blue. The mesh of blue, green, and magenta indicate the positions of α-ketoglutarate and ATP at the predicted binding sites, respectively. White and orange stick models indicate the citrate and CoA in the reported crystal structure, respectively.

## Notes

### Competing Interest Statement

The authors have declared no competing interest.

